# Zbtb11 interacts with Otx2 and patterns the anterior neuroectoderm in *Xenopus*

**DOI:** 10.1101/2023.10.20.563376

**Authors:** Yumeko Satou-Kobayashi, Shuji Takahashi, Yoshikazu Haramoto, Makoto Asashima, Masanori Taira

## Abstract

The *zinc finger and BTB domain-containing 11* gene (*zbtb11*) is expressed in the *Xenopus* anterior neuroectoderm, but the molecular nature of the Zbtb11 protein during embryonic development remains to be elucidated. Here, we show the role of Zbtb11 in anterior patterning of the neuroectoderm and the cooperative action with the transcription factor Otx2. Both overexpression and knockdown of *zbtb11* caused similar phenotypes: expanded expression of the posterior gene *gbx2* in the neural plate, and later microcephaly with reduced eyes, suggesting that a proper level of *zbtb11* expression is necessary for normal patterning of the neuroectoderm, including eye formation. Co-immunoprecipitation assays showed that Zbtb11 formed a complex with itself and with a phosphomimetic and repressive form of Otx2, suggesting that Zbtb11 forms a dimer or oligomer and interacts with Otx2 in a phosphorylation-dependent manner. Reporter analysis further showed that Zbtb11 enhanced the activity of the phosphomimetic Otx2 to repress a silencer element of the posterior gene *meis3*. These data suggest that Zbtb11 coordinates with phosphorylated Otx2 to specify the anterior neuroectoderm by repressing posterior genes.

## Introduction

During vertebrate development, the central nervous system (CNS) is derived from the neuroectoderm, which is induced and patterned by signalling molecules released from the Spemann-Mangold organizer [1,2]. The neuroectoderm forms the neural plate, and subsequently the neural tube by folding the neural plate. Before neural tube closure, the anterior portion of the neural tube is divided into three primary brain domains along the anteroposterior axis: the forebrain, midbrain, and hindbrain [3,4]. Studies of regulatory factors that play important roles in regional specification within the developing brain contribute to the understanding of how the structure and morphology of the complex brain is formed.

Using expression pattern screening with *Xenopus laevis* neurula embryos and an anterior neuroectoderm (ANE) cDNA library, we previously identified 25 uncharacterized genes [5]. Among them, we focused on an expressed sequence tag (EST) clone N21D4, because this gene is specifically expressed in the ANE, including the eye field [5]. Then, N21D4 turned out to be a gene encoding a member of the zinc finger (Znf) and Broad-complex, Tramtrack, and Bric-a-brac (BTB) domain-containing protein, Zbtb11. Zbtb family proteins contain the N-terminal BTB domain and C-terminal C2H2 type Znf domains and generally function as transcriptional repressors and chromatin silencing factors [6–8]. The BTB domain is required for the oligomerization of Zbtb proteins [9,10] and repressor functions via the recruitment of transcriptional co-repressors, such as Sin3A (SIN3 transcription regulator family member A), N-CoR (nuclear receptor corepressor 1) and SMRT (silencing mediator of retinoic acid and thyroid hormone receptor) [11–14]. The Znf domains are required for DNA binding in two different modes: direct binding to methylated CpG dinucleotides and sequence-specific DNA sites [15–17], and indirect binding through interactions with other transcription factors [18]. To date, *zbtb11* is reportedly involved in neutrophil development by repressing the expression of the tumour suppressor gene *p53* in zebrafish [19]. In addition, *zbtb11* is involved in brain development [19], but little is known about molecular features of *zbtb11* in early vertebrate development.

One of the key factors in early neural patterning is the homeodomain transcription factor Otx2. In the neuroectoderm, *otx2* is expressed in the ANE, which forms the forebrain and midbrain, and plays a critical role in the specification of the anterior portion of the head [20–24]. At the neurula stage, Otx2 represses the expression of the posterior gene *gbx2* and conversely, Gbx2 represses the expression of *otx2* in the posterior neuroectoderm [25–27]. The mutual repressive interactions between Otx2 and Gbx2 are required for the segregation of the different domains and the establishment of the midbrain and hindbrain boundary (MHB), which positions the expression of the homeobox genes *engrailed-2* (*en2)* and *pax2* [28,29] as well as *hes7.1* (*XHR1*) [30,31]. In the ANE, Otx2 initiates the gene cascade of eye field specification, including *rax* and *pax6* at the late gastrula stages, and then these eye field-specific transcription factors repress *otx2* expression to segregate the eye field from the diencephalon [32–35]. Thus, *otx2* plays an important role in the patterning of the neuroectoderm and eye field specification through regulatory interactions with other region-specific genes. To date, several reports have shown negative and positive regulation of Otx2 towards its direct target genes. For example, Otx2 interacts with the co-repressor TLE/Groucho to repress posterior genes [36,37] and cooperates with the transcription factor Sox2 to activate *rax* gene expression [38]. Furthermore, genome-wide target analysis of Otx2 in the head organizer underlying the ANE using *Xenopus tropicalis* gastrula embryos has shown that Otx2 activates head organizer genes in cooperation with the transcriptional activator Lim1/Lhx1, whereas it inhibits trunk organizer genes in cooperation with the transcriptional repressor Goosecoid (Gsc) and Tle1 [39]. In addition, our previous study suggests that Otx2 activity to repress posterior genes, *gbx2* and *meis3,* in the ANE is regulated by phosphorylation around the repression domain of Otx2 [40]. Thus, it has been shown that both transactivation and transrepression activities of Otx2 are regulated by coordinated action with binding partners and post-translational modifications.

Here, we demonstrated the biological function of Zbtb11 in early neural development using an allotetraploid species *X. laevis* and a diploid species *X. tropicalis* for gain-of- and loss-of-function analyses, respectively. Notably, both overexpression and knockdown of *zbtb11* gave rise to a similar phenotype showing posteriorization of the neuroectoderm as well as eye defects. These data suggest that Zbtb11 is required for its proper functions for anterior patterning of the neuroectoderm and eye formation in a stoichiometric manner. Furthermore, biochemical and reporter analyses showed that Zbtb11 physically interacts with Otx2 and enhances the repression activity of Otx2 in a phosphorylation-dependent manner. Together, these data provide the insights into the molecular mechanisms by which Zbtb11 functions as a regulatory component of the Otx2 repression complex for anterior neural patterning, and probably also in eye formation.

## Materials and methods

### cDNA cloning, sequence analysis and constructs

EST clones of *zbtb11* (clone name: N21D4) in *X. laevis* was previously reported [5]. The 5’-portion of *zbtb11* (2078 bp) was purchased from Open Biosystems (IMAGE 5065565) and cloned into the 3’-portion of the *zbtb11* (cloned by Dr. H. Mamada) at BamHI/AflII sites of pCSf107-mT vector [41] to reconstruct the full-length cDNA *zbtb11*.S (accession number XM_018249724). The coding sequences (CDSs) of *zbtb11.S* [amino acid numbers 1-1118] were PCR-cloned into pCSf107_Venus_mT [for Venus (an eGFP variant) constructs, gifted by Dr. N. Sudou], pCSf107_MTmT (Myc constructs), and pCSf107_HAmT (HA constructs) vectors [42]. PCR fragments for the N-terminal region including BTB domain [1–580] and C-terminal Znf domains [622–1118]) of Zbtb11.S were cloned into pCSf107-Venus_mT and pCSf107-Venus-NLS_mT. Predicted protein domains were searched using InterPro (https://www.ebi.ac.uk/interpro/). All constructs used in this study are listed in S1 Table.

### Manipulation of *Xenopus* embryos

Animal experiments were approved by the Animal Experimentation Committee at Teikyo University (approval number: 20-022), National Institute of Advanced Industrial Science and Technology (approval number: A2023-0237), and the Animal Care and Use Committees in the University of Tokyo. All animal experiments were conducted in accordance with relevant guidelines. *X. laevis* and *X. tropicalis* embryos were artificially fertilized, dejellied, and cultured in 0.1x Steinberg’s solution [43]. Embryos were staged according to the Nieuwkoop and Faber normal table [44].

### Whole-mount in situ hybridization (WISH)

DIG-labeled antisense *zbtb11* probe was transcribed with T7 RNA polymerase from *Bam*HI-linearized pCSf107-Zbtb11-T plasmid, and other antisense RNA probes were obtained as described in S2 Table. WISH assay of *X. laevis* and *X. tropicalis* embryos was performed according to the method described by Harland [45] with BM purple (Roche).

### Microinjection of mRNA and antisense morpholino oligo (MO)

Microinjection experiments were performed as previously described [40]. For mRNA synthesis, pCSf107mT constructs were transcribed using SP6 polymerase (mMESSAGE mMACHINE SP6, Thermo Fisher Scientific). Synthesized mRNAs were injected into 2- or 4-cell stage *X. laevis* embryos. Amount of injected mRNA (pg/embryo): *Venus-NLS*, 500; *Venus-Zbtb11*, 2500; *Venus-BTB*, 1500; *Venus-Znf*, 1500; *Venus-NLS-BTB*, 1500. Nuclear *β-galactosidase* (*nβ-gal*) mRNA (100 pg/embryo) was co-injected for lineage tracing. For MO injection, antisense MO against sequences near the start codon of *zbtb11* in *X. tropicalis* was obtained from Gene Tools: *zbtb11*-MO, 5’-ccaggtagctctcctcgttaga<u>cat</u>-3’ (antisense ATG codons are underlined). Standard control MO (control-MO) targeting a *β-globin* intron was used as a negative control: 5’-cctcttacctcagttacaatttata-3’. MO was dissolved in water and injected into 4-cell stage *X. tropicalis* embryos. Amount of MO (pmol/embryo): 0.5 or 1. Fluorescein isothiocyanate (FITC)-dextran (5 ng/embryo) was co-injected for lineage tracing in MO-injected experiments. Injected embryos were fixed with MEMFA (0.1 M MOPS, pH 7.4, 4 mM EGTA, 1 mM MgSO_4_, and 3.7% formaldehyde) for direct fluorescence observation or WISH. For rescue experiments, *X. tropicalis* embryos were first injected with MO and FITC-dextran into the animal pole region of one dorsal blastomere at the 4-cell stage, and then injected with *Venus-zbtb11* and *nβ-gal* mRNA into the center of one dorsoanimal blastomere on the same side at the 8-cell stage. Injected embryos were fixed at stages 38–42 with MEMFA and subjected to WISH analysis for *en2* expression. Experiments were repeated with at least two clutches.

### Fluorescence observation

mRNA for Venus-Zbtb11 constructs or Venus-NLS was injected into one blastomere at the 4-cell stage. Antisense MO for *zbtb11* (*zbtb11*-MO) or control-MO was co-injected with FITC-dextran as a tracer into one blastomere at the 4-cell stage. Injected embryos were incubated until embryos reached the appropriate stages, and fixed in MEMFA for 1 or 1.5 hours. Expression of Venus-Zbtb11 constructs and Venus-NLS was observed using an LSM 710 confocal microscope (Zeiss) and a fluorescent microscope (Leica). Expression of FITC-dextran in MO-expressing embryos was observed using a fluorescent microscope (Leica).

### Measurements of the eye-size, lengths from the cement gland to the eye, and body lengths

Injected embryos for phenotypic analyses were fixed at tailbud stages with MEMFA for 1 or 1.5 hours. The area of the eye vesicle, the distance from the cement gland to the center of the eye, and the rostrocaudal length of the body axis were measured using the Fiji software [46]. Two-tailed Student’s t-test was used after a one-way analysis of variance (ANOVA) to calculate the statistical significance. For rescue experiments, a relative position of *en2* at the MHB on the MO-injected side was measured from that on its uninjected side as the reference point (0 μm); the anterior or posterior shift of *en2* expression on MO-injected side was measured as + or - μm distance using Fiji [46].

### Western blotting

mRNA was injected into both blastomeres in the animal pole region at the 2-cell stage and embryonic lysates were prepared at the appropriate stages. SDS-polyacrylamide gel electrophoresis (PAGE) was carried out with 7.5-12.5% polyacrylamide gels, and western blotting was performed as described elsewhere [42] using anti-GFP IRDye 800 conjugated (Rockland Immunochemicals Inc., 600-132-215, 1:5000 dilution), anti-β-tubulin (Sigma, T4026, 1:5000), anti-Myc (9E10, 1:10000) and anti-HA (12CA5, 1:10000) antibodies. Protein bands were detected using Odyssey Infrared Imaging System (LI-COR, ODY-9201). To determine the relative expression levels of Venus-Zbtb11 and Venus-NLS, the band intensity of Venus fused protein was divided by that of *β*-tubulin as a loading control.

### Co-immunoprecipitation (Co-IP) assay

mRNA was injected into both blastomeres at the 2-cell stage, and embryonic lysates were prepared at gastrula stages (stages 10.5-11). Co-IP assays were performed as described elsewhere [42] with minor modifications. Embryonic lysates were incubated with the appropriate antibody for 2 hours at 4℃, and subsequently incubated with 20 μL of protein G-agarose beads (Roche) or protein A-agarose beads (Sigma) for 2-3 hours at 4℃. Bound proteins were eluted by boiling in SDS sample buffer, separated by SDS-PAGE (7.5% or 10% gel), and analyzed by western blotting.

### Luciferase reporter analysis

SOP-FLASH [38] or pGL4p-*meis3*-D2-luc reporter DNA [40] was co-injected with mRNA into two dorsal animal blastomeres at the 4-cell stage or into both animal blastomeres at the 2-cell stage. Five pools of three injected embryos were assayed for luciferase activity at stage 12 for SOP-FLASH, and at stage 10.5 for pGL4p-*meis3*-D2-luc reporter. The luciferase reporter assay system (Promega) was used according to the manufacturer’s protocol. Two-tailed Student’s t-test was used after a one-way analysis of variance (ANOVA) to calculate the statistical significance. Experiments were repeated at least three times except for the luciferase assay using Otx2 phosphomutants, and a representative result is exhibited when similar results were obtained.

## Results

### Expression profile of *zbtb11*

The allotetraploid *X. laevis* has two homeologs of *zbtb11*, *zbtb11.L* and *zbtb11.S*. The previously isolated EST clone N21D4 [5] was identified as a partial sequence of *zbtb11.S*. The full-length cDNA clone of *zbtb11.S* (3.4 kb) was reconstructed using PCR clones and an IMAGE clone, which overlapped with the nucleotide sequence of N21D4, as described in Materials and Methods. We first compared developmental expression between *zbtb11.L*, *zbtb11.S*, and *X. tropicalis zbtb11* using published RNA-sequencing (RNA-seq) datasets [47,48]. As shown in S1A Fig, *zbtb11.S* showed much higher expression levels than *zbtb11.L* at oocyte stages I-VI and likewise higher maternal expression levels from eggs to stage 8. Note that the apparent gradual increase of maternal expression owes to polyadenylation of mRNA after fertilization. From stage 8 onwards, though *zbtb11.S* showed higher expression levels than *zbtb11.L* until the mid-neurula stage (stage 15), both genes showed similar zygotic expression patterns with peaks at stages 9 and 15. *X. tropicalis zbtb11* showed maternal and zygotic expression patterns similar to *X. laevis* until neurula stages (S1B Fig). However, in contrast to *X. laevis*, its expression increased after around stage 30. These data suggest that temporal expression patterns of *zbtb11* genes until late neurula stages appear to be evolutionally conserved between the *X*. *laevis* homeologs and *X*. *tropicalis* ortholog. In this study, we used *zbtb11.S* constructs for expression and functional studies.

Spatiotemporal expression of *zbtb11* was examined to predict its function in embryonic development using whole-mount in situ hybridization (WISH) of *X. laevis* embryos with *zbtb11.S* as probe, which probably crossreacted with *zbtb11.L* (91.06% identity in the CDS). As shown in Fig 1, *zbtb11* was broadly expressed in the animal hemisphere at the 4-cell and early blastula stage (stage 8). Then, *zbtb11* was detected in the dorsal mesoderm (dm) at the mid-gastrula stage (stage 11), in the sensorial layer of the neuroectoderm (sl) at the late gastrula stage (stage 12.5), and was gradually restricted to the eye field (ef) at the early neurula stage (stage 14). Subsequently, *zbtb11* expression was restricted to the presumptive eyes (pey) and brain (br) (stage 23), and the forebrain (fb) and midbrain (mb), eyes (ey), otic vesicles (ov), branchial arches (ba), a posterior part of the hindbrain (ph), somites (sm) and presomitic mesoderm (psm) (stages 35-36). Note that *zbtb11* was highly expressed at the midbrain (stages 35-36), where *otx2* is expressed. These data suggest that *zbtb11* exerts pleiotropic roles in various organs and tissues including the neuroectoderm and the eyes.

**Fig 1.**
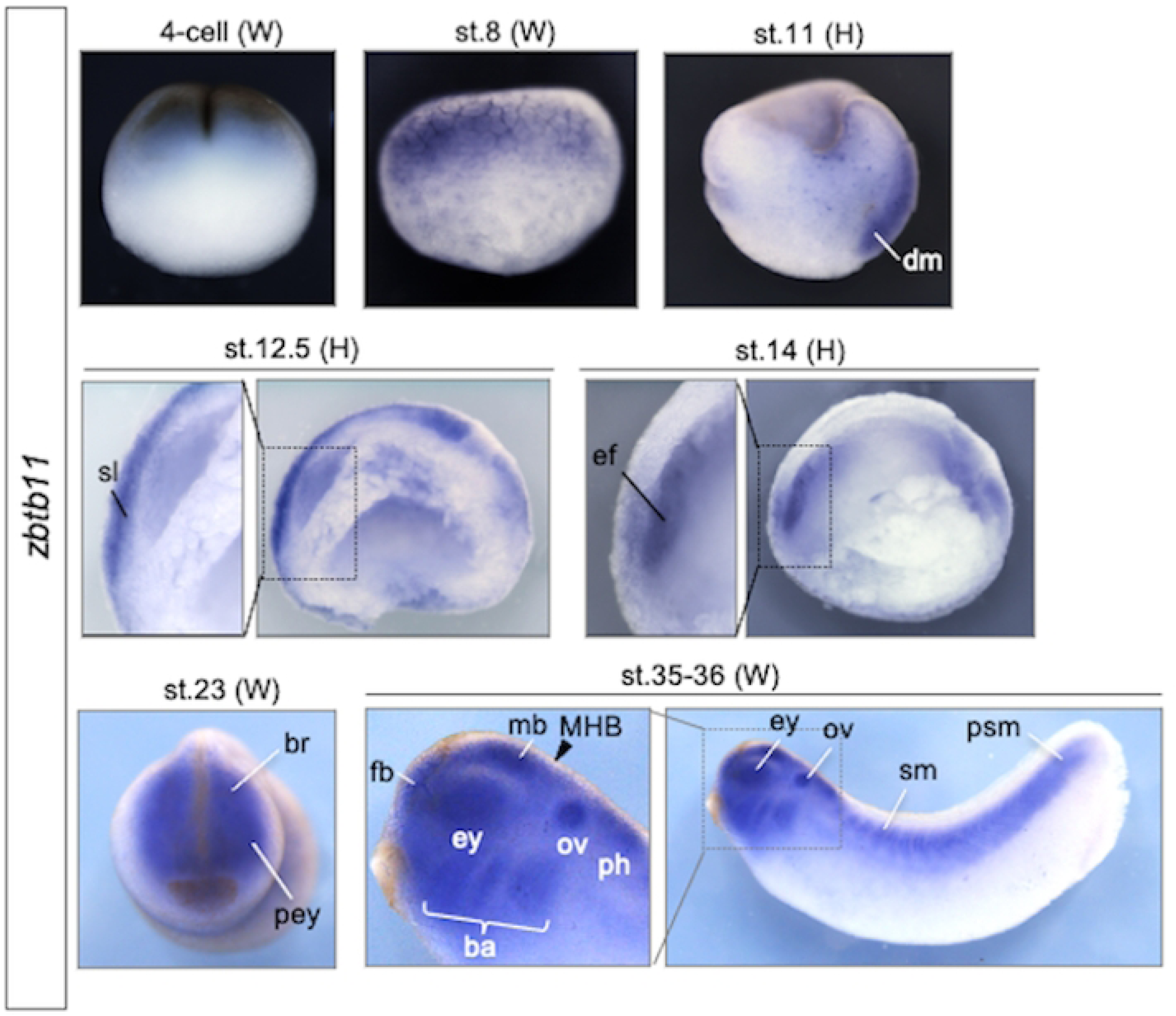
Developmental expression patterns of zinc finger and BTB domain-containing 11 (zbtb11). Whole-mount in situ hybridization (WISH) analysis of *zbtb11*. Stages are indicated at the upper of each panel. Whole embryo (W) and hemisection (H) are as indicated. Lateral view with animal pole side up (4-cell, st. 8 and 11). Lateral view with the dorsal side up (st.12.5, 14, 35-36). Anterior view with the dorsal side up (st. 23). Dashed boxes indicate enlarged images of st. 12.5, 14 and 35-36. An arrowhead indicates the position of the midbrain and hindbrain boundary (MHB), and a bracket indicates *zbtb11* expression in the branchial arches. ba, branchial arches; br, brain; dm, dorsal mesoderm; ef, eye field; ey, eyes; fb, forebrain; mb, midbrain; MHB, midbrain and hindbrain boundary; ov, otic vesicles; pey, presumptive eyes; ph, posterior part of the hindbrain; psm, presomitic mesoderm; sl, sensorial layer; sm, somites.

### Molecular features of Zbtb11

Amino acid sequence alignment of Zbtb11 among different vertebrate species showed that Zbtb11 had a conserved region (CR1, light blue box) including the integrase-like histidine-histidine-cysteine-cysteine motif (HHCC, purple box) and BTB domain (brown box) on the N-terminal side, 12 C2H2 type Znf domains (pink boxes) and conserved regions (CR2 and CR3, light blue boxes) on the C-terminal side (S2 Fig). To examine the region responsible for the activity of Zbtb11, we divided full-length Zbtb11 [amino acid positions (aa) 1–1118] into two parts: (i) the N-terminal CR1 region containing the HHCC motif and BTB domain [aa 1–580, named BTB], and (ii) the C-terminal region containing Znf domains, CR2 and CR3 [aa 622–1118, Znf]. We first checked the stability of Zbtb11 protein exogenously expressed in the embryos. Western blotting analysis of Myc-tagged Zbtb11 (Myc-Zbtb11) showed many bands of degradation products lower than the band of the full-length protein (S3A Fig). Compared to this, Venus-fused Zbtb11 (Venus-Zbtb11) appeared to be more stable, because band intensities of its degradation products were lower than those of Myc-Zbtb11 (S3A and S3B Fig; compare bands with magenta dots). Regarding Zbtb11 deletion constructs, Venus-BTB degraded in a fashion similar to Venus-Zbtb11 (compare bands with two magenta dots and blue dots), whereas Venus-Znf appeared to be more stable than Venus-BTB (S3B Fig). This data implies that the proteolytic degradation of Zbtb11 mainly occurred in the CR1 region including the BTB domain and the following linker sequence.

We next compared the expression level of Venus-Zbtb11 with that of Venus-NLS during early development (S4 Fig). The fluorescent signal of Venus-Zbtb11 was weaker than that of Venus-NLS at the early neurula stage (stage 14) and was reduced at later stages, whereas the fluorescent signal of Venus-NLS was maintained until tailbud stages (stages 28–30) (S4A Fig). Consistent with this, western blotting analysis showed that protein levels of Venus-Zbtb11 were much lower than those of Venus-NLS during early development (S4B and S4C Fig). This data also supports the possibility that Zbtb11 is an unstable protein.

We next examined the subcellular localisation of Zbtb11 in *Xenopus* embryos using Venus-Zbtb11 and its deletion mutants (Fig 2A). The confocal microscopic observation of Venus fluorescence showed that Venus-Zbtb11 was localised in the nucleus (Fig 2B), suggesting its function as a nuclear protein. Deletion analysis showed that Venus-BTB was mainly localised in the cytoplasm, whereas Venus-Znf was localised in the nucleus (Fig 2B). Furthermore, when the nuclear localisation signal (NLS) of SV40 large T was added to Venus-BTB to construct Venus-NLS-BTB, the subcellular localisation changed from the cytoplasm to the nucleus (Fig 2B). These data indicate that the NLS of Zbtb11 resides in the C-terminal region including Znf domains.

**Fig 2.**
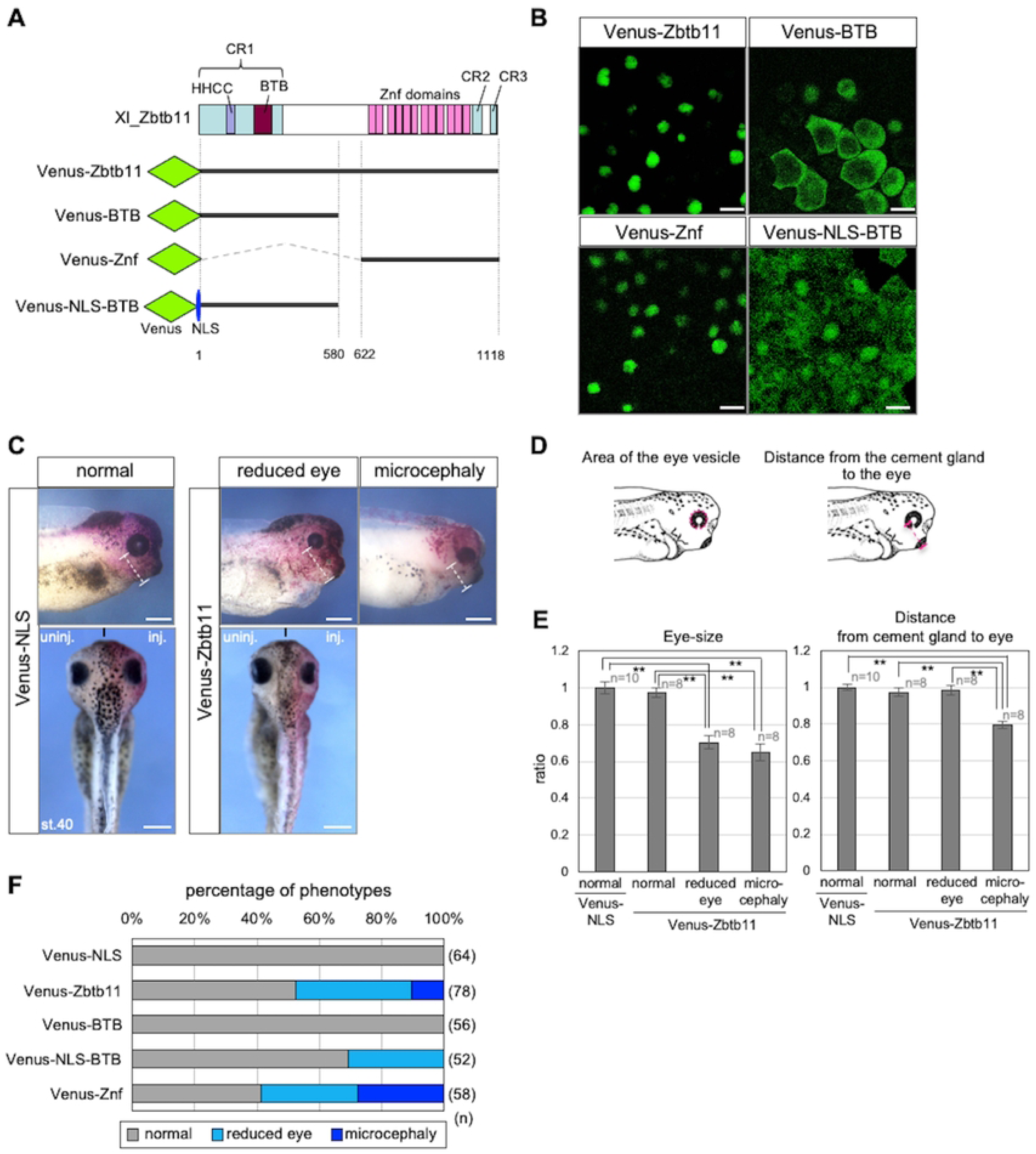
Subcellular localisation and activities of full-length Zbtb11 or its deletion constructs in eye development. (A) Schematic protein structure of *Xenopus laevis* (Xl) Zbtb11 and the representation of Venus-fused constructs of Zbtb11. The N-terminal conserved region (CR1, light blue box) including the integrase-like histidine-histidine-cysteine-cysteine motif (HHCC, purple box) and BTB domain (brown box), C2H2 type Znf domains (pink boxes), and the C-terminal conserved regions (CR2 and CR3, light blue boxes) are indicated. The N-terminal Venus tag is indicated as green rhombuses, and the nuclear localisation signal (NLS) is indicated as a blue ellipse. The regions of Zbtb11 constructs are indicated by thick lines and the positions of amino acid residues are indicated. (B) Subcellular localisation of Zbtb11. mRNA for Venus-fused Zbtb11 constructs was injected at the 4-cell stage and detected by confocal microscopy at the gastrula stage. Scale bars: 25 μm. (C-F) Morphological changes by overexpression of Zbtb11 and its deletion constructs. mRNA for each Venus-Zbtb11 construct or Venus-NLS was co-injected with *nβ-gal* mRNA as a tracer into one dorsal blastomere at the 4-cell stage, and phenotypes were checked at the tailbud stages. The amount of injected mRNA was adjusted to obtain equal number of moles. (C) Representative images of phenotypes at stage 40 (st.40). Phenotypes are categorised normal-looking (normal), reduced eyes (reduced eye), and small head with ventrally positioned eyes (microcephaly). White dashed lines indicate the length from the cement gland to the center of the eye. Anterior to the right, dorsal is up (upper panels). Dorsal view with the anterior side up (lower panels). Black short lines into the lower panels indicate the midline of the embryo. inj., injected side; uninj., uninjected side. Scale bars: 500 μm. (D) Schematic images of the measurements for the eye-size and the lengths from the cement gland to the eye. The eye vesicle was approximated by ellipse and the area of the eye vesicle was measured. The ratio of eye-size of tested samples was calculated by dividing with the area of the eye vesicle in Venus-NLS-expressing embryos (left). The length from most distal part of the cement gland to the center of the eye was measured. The ratio of length from the cement gland to the eye of tested samples was calculated by dividing with those in Venus-NLS-expressing embryos (right). Magenta dashed circle, the eye vesicle; magenta dashed line, the length from the cement gland to the center of the eye. Images of tailbud stage embryos are adapted from Nieuwkoop and Faber, 1994. (E) Quantitative analysis of the eye-size and the length from the cement gland to the eye. \*\**P*<0.01 (Student’s t-test); error bars, s.e.m.. (F) Occurrence rates of phenotypes. Color code of bars represents each phenotype in lower panel. Experiments were repeated with at least three clutches. n, the total number of samples (E,F).

### Overexpression of Zbtb11 leads to eye defects and posteriorization of the anterior neural plate

For gain-of-function experiments, mRNA for Venus-Zbtb11 or Venus-NLS as a negative control was co-injected with *nβ-gal* mRNA as a tracer into one dorsal-right blastomere at the 4-cell stage, aiming for expression in the ANE, and phenotypes were scored at stages 35–40. The amount of injected mRNA for Venus-Zbtb11 ranged from 625 pg to 2.5 ng/embryo, since at least 2.5 ng mRNA was required to cause any phenotypic abnormalities. As shown in Fig 2C, overexpression of Venus-Zbtb11 exhibited eye and brain defects in a certain percentage of the injected embryos, which could be categorised into either ‘reduced eyes’ or a small head with ventrally positioned small eyes (‘microcephaly’), compared to the uninjected left side and uninjected embryos. To quantitatively evaluate morphological defects, we measured the area of the eye vesicle and the distance from the cement gland to the eye (Fig 2D). As shown in Fig 2E, there was no significant difference in eye size between Venus-Zbtb11-expressing embryos with the normal-looking phenotype and Venus-NLS-expressing embryos. By contrast, the eye size in Venus-Zbtb11-expressing embryos with the reduced eye phenotype or microcephaly was significantly smaller than those in Venus-NLS-expressing embryos (Fig 2E left graph; 0.65 to 0.70-fold decrease in average). Furthermore, the distance from the cement gland to the eye in Venus-Zbtb11-expressing embryos with microcephaly was significantly shorter than those in Venus-Zbtb11-expressing embryos with the normal-looking and reduced eye phenotype or Venus-NLS-expressing control embryos (Fig 2E right graph; 0.80-fold decrease in average). Note that about half of the Zbtb11-overexpressing embryos did not exhibit any phenotypic defects (Fig 2F). This might be due to its low protein stability, as suggested above (S3 and S4 Figs). To examine which regions of Zbtb11 cause reduced eye size or microcephaly, Venus-NLS-BTB and Venus-Znf was compared. The data showed that Venus-Znf caused both phenotypes as does the full-length construct (Fig 2F), indicating that the C-terminal region containing Znf domains seemed to be sufficient for exhibiting the phenotypes by Zbtb11. In addition, Venus-NLS-BTB also caused the reduced eye phenotype, but not microcephaly, implying that eye defects result from both effects of the N-terminal region containing the BTB domain and the C-terminal region containing Znf domains. Note that Venus-BTB did not cause any abnormalities (Fig 2F), suggesting that nuclear localisation is necessary for the activity of the N-terminal region containing the BTB domain.

To examine how Zbtb11 overexpression results in the reduced eye phenotype and microcephaly, WISH analysis was performed for injected embryos at the late gastrula (stages 12–12.5) and early neural stages (stages 13–14). We first tested the effect of exogenous Zbtb11 on anteroposterior patterning of the neuroectoderm by focusing on the regional markers *otx2*, *gbx2*, and *pax2* (Figs 3A-L). Venus-Zbtb11 and -Znf slightly reduced the expression of *otx2* at the posterior end (green arrowheads; that is, anterior reduction) and anteriorly expanded the *otx2*-expressing domain (magenta arrowheads) (Fig 3B and 3C). In addition, Venus-Zbtb11 and -Znf caused the anterior expansion of *gbx2* expression (Fig 3F and 3G; magenta arrowheads; that is, posterior expansion) and the disappearance of *pax2* expression at the MHB (Fig 3J and 3K; arrows). By contrast, Venus-NLS-BTB barely affected expression of *otx2*, *gbx2* and *pax*2 (Fig 3D, 3H and 3L), compared to that of the Venus-NLS control (Fig 3A, 3E and 3I) or the uninjected side (Fig 3D, 3H and 3L). These data suggest that overexpression of Zbtb11 promotes posteriorization of the ANE through its C-terminal region containing Znf domains. We next examined expression of the eye-field specific genes, *rax* and *pax6* at the late gastrula (Figs 3M-T). As expected from the reduced eye phenotype, Venus-Zbtb11 and -Znf reduced *rax* and *pax6* expression on the injected side (Fig 3N, 3O, 3R, and 3S; green arrowheads; *pax6* expression in the somites (sm) was not affected), which might be due to the reduction of *otx2* expression and the expansion of *gbx2* (see Fig 3B, 3C, 3F, and 3G). By contrast, Venus-NLS-BTB did not largely affect *rax* expression (Fig 3P), but reduced *pax6* expression in the eye field (Fig 3T; green arrowhead). Notably, Venus-Zbtb11 and Venus-NLS-BTB ectopically induced *pax6* expression in the region anterior to the eye field (Fig 3R, 3R’, 3T and 3T’), unlike Venus-Znf (Fig 3S and 3S’). Because Venus-NLS-BTB did not affect anterior patterning (Fig 3D, 3H, and 3L) or *rax* expression (Fig 3P), it is possible that the effect of the N-terminal BTB region on *pax6* expression is direct and context-dependent. These data suggest that Zbtb11 is involved in both anterior patterning of the neuroectoderm and eye formation through the Znf domains and the N-terminal BTB region, respectively.

**Fig 3.**
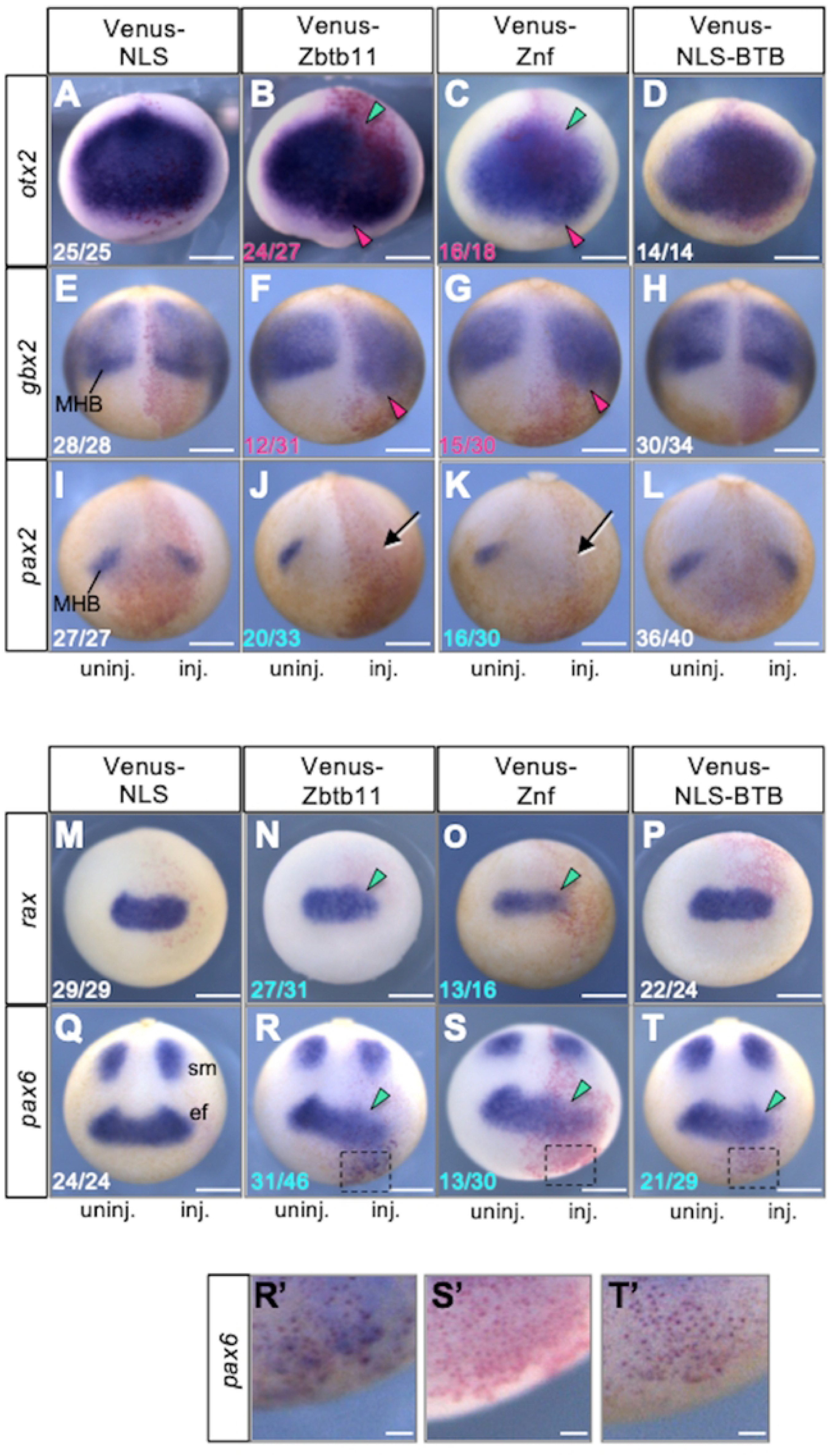
Effects of Zbtb11 and its deletion constructs on neural marker genes. Effects of overexpression of Zbtb11 and its deletion constructs on anteroposterior patterning of the neuroectoderm (A-L) and early eye formation (M-T). mRNA for each Venus-Zbtb11 construct or Venus-NLS was co-injected with *nβ-gal* mRNA as a tracer into one dorsal blastomere at the 4-cell stages. WISH analysis of *otx2* (A-D), *gbx2* (E-H), *pax2* (I-L), *rax* (M-P), and *pax6* (Q-T) was performed at the late gastrula stage (stages 12–12.5) and early neurula stage (stages 13–14). Experiments were carried out with at least three clutches. Expression of marker genes was compared between injected (inj.) and uninjected areas (uninj., as a negative control). Fractional numbers indicate the numbers of the embryos presenting the phenotype per scored embryos (numbers in white, no or subtle effects on gene expression; magenta, expanded expression; blue, reduced expression). Colored arrowheads: green, reduction of gene expression; magenta, anterior expansion. Arrows, the absence or strong reduction of *pax2* expression. Anterior view with the dorsal side up (*otx2* and *rax*), dorsal view with the posterior side up (*gbx2* and *pax2*), and dorsoanterior view with the posterior side up (*pax6*). ef, eye field; MHB, the midbrain and hindbrain boundary; sm, somites. (R’-T’) Enlarged images of R-T (black dashed boxes). Scale bars: 500 μm in A-T, 100 μm in R’-T’.

### Knockdown of *zbtb11* causes posteriorization similar to the overexpression phenotype

To investigate the involvement of *zbtb11* during embryogenesis, we carried out knockdown experiments using *X. tropicalis* embryos by injecting antisense morpholino oligos (MOs) for *X. tropicalis zbtb11* targeting its 5ʹ portion, including the start codon (*zbtb11*-MO). Because *X. tropicalis* is a diploid species closely related to the allotetraploid species *X. laevis* [47,48], it is suitable for MO knockdown experiments. We first checked the specificity of *zbtb11*-MO using a construct of the *eGFP* CDS possessing the MO-target sequence at the N-terminus, named Zbtb11-ATG-eGFP. *zbtb11-ATG-eGFP* mRNA was first injected into both blastomeres at the 2-cell stage, and then *zbtb11-* MO or control-MO was injected into all blastomeres at the 4-cell stage. Injected embryos were subjected to western blotting at the gastrula stage. The eGFP band translated from *zbtb11-ATG-eGFP* mRNA was diminished by *zbtb11*-MO, but not by the control-MO, compared to that of the uninjected control (S5 Fig, upper panel). In addition, expression levels of the loading control, β-tubulin, were not affected by any MOs (S5 Fig, lower panel). These data suggest that *zbtb11*-MO specifically inhibits the translation of *zbtb11* mRNA.

We then injected *zbtb11*-MO or control-MO with FITC-dextran as a tracer into the dorsoanimal region of one blastomere at the 4-cell stage, and phenotypes of injected embryos were scored at tailbud stages (stages 40–43). Compared to control-MO-injected embryos, *zbtb11*-MO injection resulted in two abnormal phenotypes: normal body size with reduced forebrain and eye (microcephaly) (Fig 4A) or shortened body length with microcephaly (short axis) (Fig 4B), in a dose-dependent manner (Fig 4C). Quantitative analysis showed that the eye size in *zbtb11* morphants with microcephaly was significantly smaller (Fig 4D left; 0.45 fold in average), and the length of the body size of the morphants with short-axis was significantly shorter than those of control-MO-injecting embryos (Fig 4D right; 0.81 fold in average).

**Fig 4.**
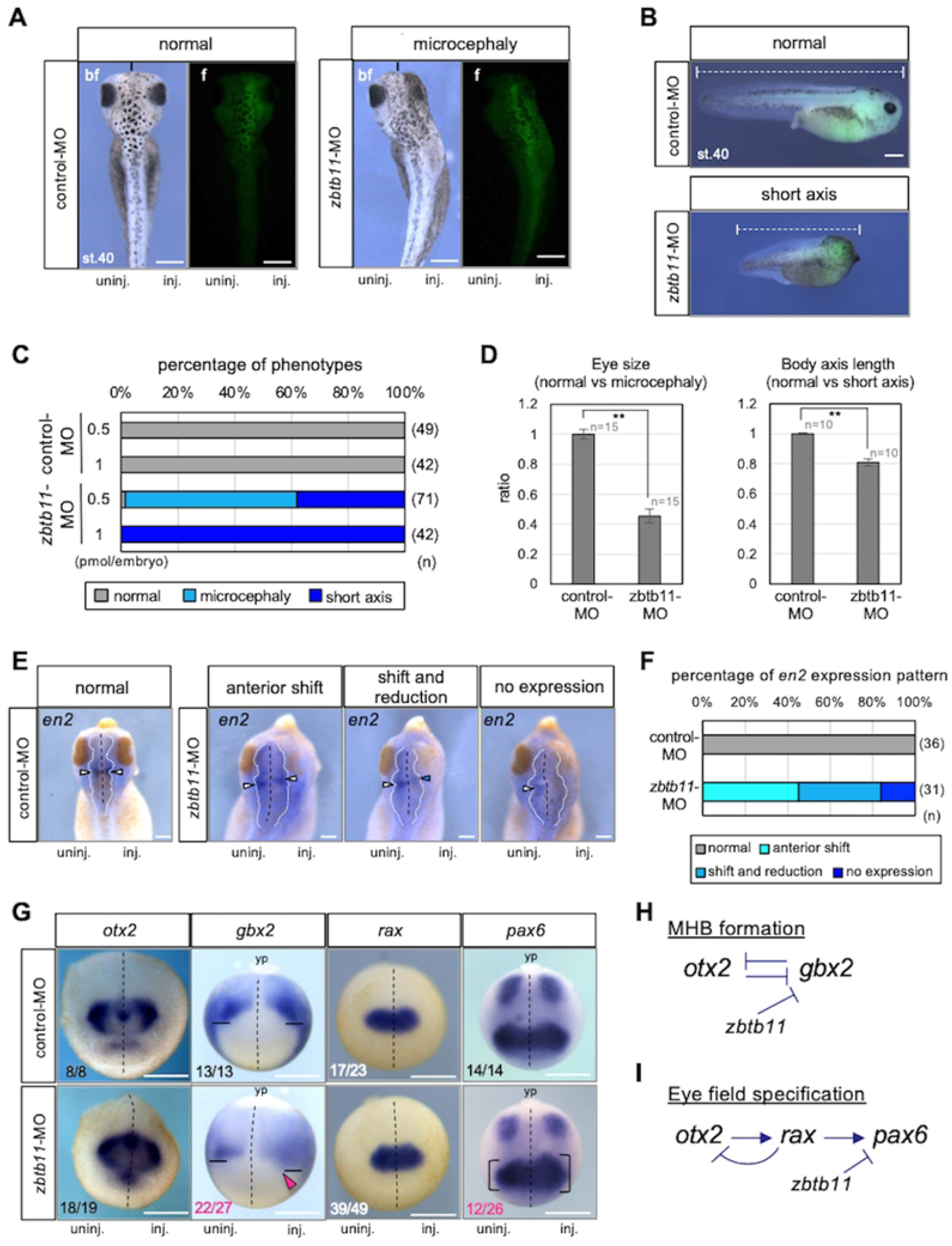
Knockdown of *zbtb11* results in brain and eye defects. For *zbtb11* knockdown experiments using morpholino oligo (MO), *zbtb11*-MO or control-MO was injected with FITC-dextran as a tracer into the animal pole region of one dorsal blastomere at the 4-cell stage. (A,B) Morphological appearances of MO-injected embryos at the tailbud stage. Representative images of normal-looking control-MO-injected embryos (normal) at stage 40 (st.40) and *zbtb11*-MO-injected embryos with microcephaly (A) or with short axis (B) are shown by bright field (bf) and fluorescent (f) images (A) or merged images of the injected right side (B). Fluorescence of FITC-dextran indicates the presence of MO. The black short lines indicate the anterior end of the midline (A). White dashed lines indicate the rostrocaudal length of the body (B). (C) Percentages of phenotypes of injected embryos presented in (A,B). Either 0.5 or 1 pmol of MO was injected per embryo as indicated. Experiments were repeated with at least three clutches. (D) Quantitative analysis of the eye-size (left) and the rostrocaudal length of the body (right). The eye vesicle was approximated as an ellipse and the area of the eye vesicle was measured. Eye sizes and body lengths were normalized using the average eye size and body length of control-MO-injected embryos. (E) WISH analysis for the midbrain-hindbrain boundary (MHB) marker *engraild 2* (*en2*) at the tailbud stage. Experiments were repeated with two clutches. Dorsal view with the anterior side up. The mode of *en2* expression is categorized: normal expression ‘normal’, ‘anterior shift’, ‘shift and reduction’, and ‘no expression’. Black dashed lines, the midline of the brain; white dotted lines, outline of the brain. Arrowheads in white, normal expression of *en2* at the MHB; those in blue, reduction of *en2* at the MHB. (F) The proportion of embryos exhibiting *en2* expression modes presented in (E). (G) Effects of *zbtb11* knockdown on anteroposterior patterning of the neuroectoderm and eye field formation. WISH analysis for *otx2*, *gbx2*, *rax and pax6* at stages 13–14. Fractions indicate the numbers of the embryos presenting the phenotype per scored embryos (numbers in white or black, minor effects on gene expression; magenta, expanded expression). Dashed line, the midline of the embryo; solid line, the anteriormost position of *gbx2* expression; magenta arrowhead, anterior expansion of *gbx2*; bracket, the size of the eye field expressing *pax6*. Anterior view with the dorsal side up (*otx2*, *rax*), and dorsal view with the posterior side up (*gbx2*, *pax6*). yp, yolk plug. inj., MO-injected side; uninj., uninjected side (A,E,G). n, the total number of each sample (C,D,F). Scale bars: 200 μm (A,B), 100 μm (E), 500 μm (G). Amounts of injected MOs (pmol/embryo): 0.5 (A,B,D-G), 0.5 or 1 (C). (H,I) Schematic models of gene interactions in MHB formation and eye-field formation. Mutual repression between *otx2* and *gbx2* (H) and the gene cascade of *otx2*, *rax*, and *pax6* (I) have well been documented (see the text). *zbtb11*-MO experiments suggest that Zbtb11 represses *gbx2* expression anterior to the MHB and represses *pax6* but not *rax* in the eye field. Arrow, activation; T mark, inhibition.

To further investigate the cause of eye and brain defects by *zbtb11* knockdown, we examined the expression of the MHB marker gene *en2* in morphants with microcephaly by WISH. Compared to control-MO, *zbtb11*-MO caused aberrant *en2* expression on the MO-injected side (Fig 4E and 4F). The mode of *en2* expression was categorized into three types compared with the MO-uninjected side: ‘anterior shift’, anterior shift with reduction (‘shift and reduction’) and ‘no expression’ (Fig 4E), and scored (Fig 4F). The data showed that more than 80% of injected embryos exhibited the anterior shift of *en2* expression by combining the phenotypes of ‘anterior shift’ and ‘shift and reduction’, suggesting that *zbtb11* knockdown causes the expansion of the hindbrain, leading to microcephaly (Fig 4A).

To examine earlier effects of *zbtb11* knockdown on the patterning of the ANE, WISH analysis of MO-injected embryos was performed at stages 13–14 for *otx2*, *gbx2*, *rax* and *pax6* at the early neurula stage (stages 13–14). As shown in Fig 4G, *otx2* expression appeared to be unaffected by *zbtb11*-MO injection, but the *gbx2* expression domain was expanded into the anterior region with disappearance of the sharp border at the MHB (magenta arrowhead), compared to that of the uninjected side. This data suggests that Zbtb11 is required for anterior boundary formation of *gbx2* expression. Furthermore, both overexpression and knockdown of *zbtb11* similarly caused microcephaly (Fig 2C and 4A) and the anterior expansion of *gbx2* (Fig 3F and 4G), implying that proper Zbtb11 expression levels are required for its normal function in the patterning of the neuroectoderm. In addition, though it has been postulated that *otx2* and *gbx2* mutually repress each other (Fig 4H), it is unclear why *gbx2* expression was anteriorly expanded by *zbtb11*-MO in spite of the existence of the clear posterior boundary of *otx2* expression. This will be discussed later. Regarding the effect of *zbtb11* knockdown on eye field marker genes, *rax* expression did not change significantly, but the *pax6* expression domain was expanded in half of *zbtb11*-morphants (12/26), compared to that of the uninjected side (Fig 4G, comparing the sizes of brackets between MO-injected and MO-uninjected sides). According to the gene cascade of eye field specification [35] (Fig 4I), this data suggests that *zbtb11* functions downstream of *rax* and upstream of *pax6*. In this case, the loss-of-function phenotype of *zbtb11* (posterior expansion of *pax6* expression) was opposite to the gain-of-function phenotype (posterior reduction of *pax6* expression) (see Figs 3R and 4G), unlike the effect on *gbx2* (see Figs 3F and 4G). These data suggest that *zbtb11* is required for proper eye formation through *pax6* in a mechanism different from anteroposterior patterning through *gbx2*.

To validate the specificity of *zbtb11*-MO, we performed mRNA rescue experiments. *zbtb11*-MO or control-MO was injected with FITC-dextran into the animal pole region of one dorsal blastomere at the 4-cell stage, and *Venus-zbtb11* mRNA at three doses (31, 63, and 125 pg/embryo) was co-injected with *nβ-gal* mRNA into the one dorsoanimal blastomere on the same side at the 8-cell stage. WISH analysis of injected embryos was performed at stages 38–42 for *en2* mRNA to test whether the anterior shift of *en2* expression in *zbtb11*-morphants is rescued by *Venus-zbtb11* mRNA (Fig 5A). The disposition of *en2* expression between the MO-injected and uninjected sides was measured as in Fig 5B. As shown in Fig 5C, *en2* expression was significantly shifted anteriorly by *zbtb11*-MO injection compared to control-MO, and this anterior shift was significantly rescued by 63 pg/embryo of *Venus-zbtb11* mRNA, but not by 31 or 125 pg/embryo. However, the rescue with mRNA was not complete, because the expression of *en2* was still shifted anteriorly compared to control-MO injected embryos (\*\**P* < 0.01). These data suggest that the specificity of *zbtb11*-MO is verified but to some extent.

**Fig 5.**
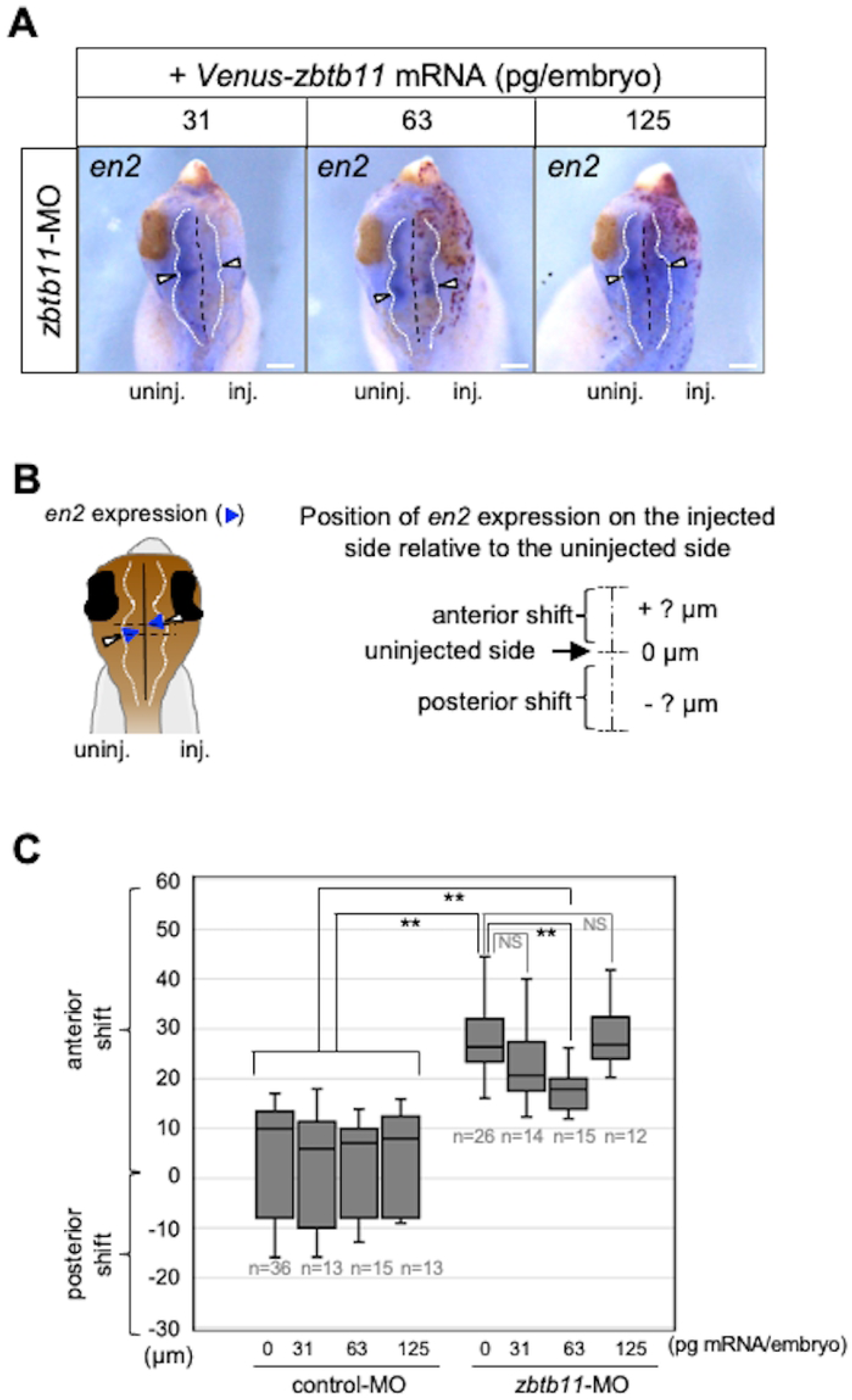
mRNA rescue experiments for the anterior shift of *en2* expression in *zbtb11*-morphants. For rescue experiments, *zbtb11*-MO or control-MO was injected into the animal pole region of the right blastomere at the 4-cell stage, and three doses of *Venus-zbtb11* mRNA (31, 63, or 125 pg/embryo) were injected into the dorsoanimal blastomere on the same side at the 8-cell stage. WISH analysis was performed for *engraild 2* (*en2*) at the tailbud stage. (A) Representative images of *en2* expression. Dorsal view of the head region with the anterior side up. Black dashed lines, the midline of the brain; white dotted lines, outline of the brain; white arrowheads, *en2* expression at the MHB. Red-gal stained cells indicate *Venus-zbtb11* mRNA expressing cells. inj., injected side; uninj., uninjected side. Scale bars: 100 μm. (B) A schematic presentation of the method to measure the position of *en2* expression on the injected side relative to the uninjected side at the MHB. Blue arrowheads stand for *en2* expression and white arrowheads indicate the middle of *en2* expression at the lateral edge. Black line, the midline of the brain; white dotted lines, outline of the brain. Perpendicular lines were drawn from the middle of the *en2* expression at the lateral edge on both the injected (inj) and uninjected (uninj) sides (dotted black lines). The disposition of *en2* expression on the injected side relative to the uninjected side was measured as the distance between these two perpendicular lines (anterior or posterior shifts in + or – μm, respectively). (C) *zbtb11* mRNA partially rescues the anterior shift of *en2* expression in *zbtb11*-morphants. The data is presented using the box and wisker plot. Amounts of co-injected *Venus-zbtb11* mRNA (0, 31, 63, or 125 pg/embryo) with control-MO or *zbtb11*-MO (0.5 pmol/embryo) are as indicated. n, the total number of each sample. \*\**P*<0.01 (Student’s t-test); error bars, s.e.m.; NS, not significant.

### Zbtb11 forms a complex with itself and Otx2

As mentioned above, gain-of-function and loss-of-function analyses suggest that proper Zbtb11 expression levels are required for its normal function in the neuroectoderm. Therefore, we hypothesised that Zbtb11 forms a protein complex in a stoichiometric manner with itself or partner proteins. We first tested the interaction of Zbtb11 with itself by co-immunoprecipitation (Co-IP) assays using full-length Zbtb11 and its deletion constructs. Embryonic lysates expressing Venus-Zbtb11 constructs with Myc-Zbtb11 were immunoprecipitated with anti-Myc antibody and subjected to western blotting with anti-GFP or anti-Myc antibody. As shown in Fig 6A, the band of Venus-Zbtb11 co-immunoprecipitated with Myc-Zbtb11 was strongly detected (magenta arrowhead) compared to the weak non-specific band seen in the absence of Myc-Zbtb11 (asterisk). This data suggests that Zbtb11 forms dimers or oligomers with itself. Furthermore, Venus-NLS-BTB and Venus-Znf clearly co-immunoprecipitated with Myc-Zbtb11, though the band of Venus-Znf was weaker than that of Venus-NLS-BTB (Fig 6A, green arrowheads), indicating that dimerization or oligomerization of Zbtb11 occurs through its N-terminal region including the BTB domain, and also through the C-terminal region including Znf domains to a lesser extent.

**Fig 6.**
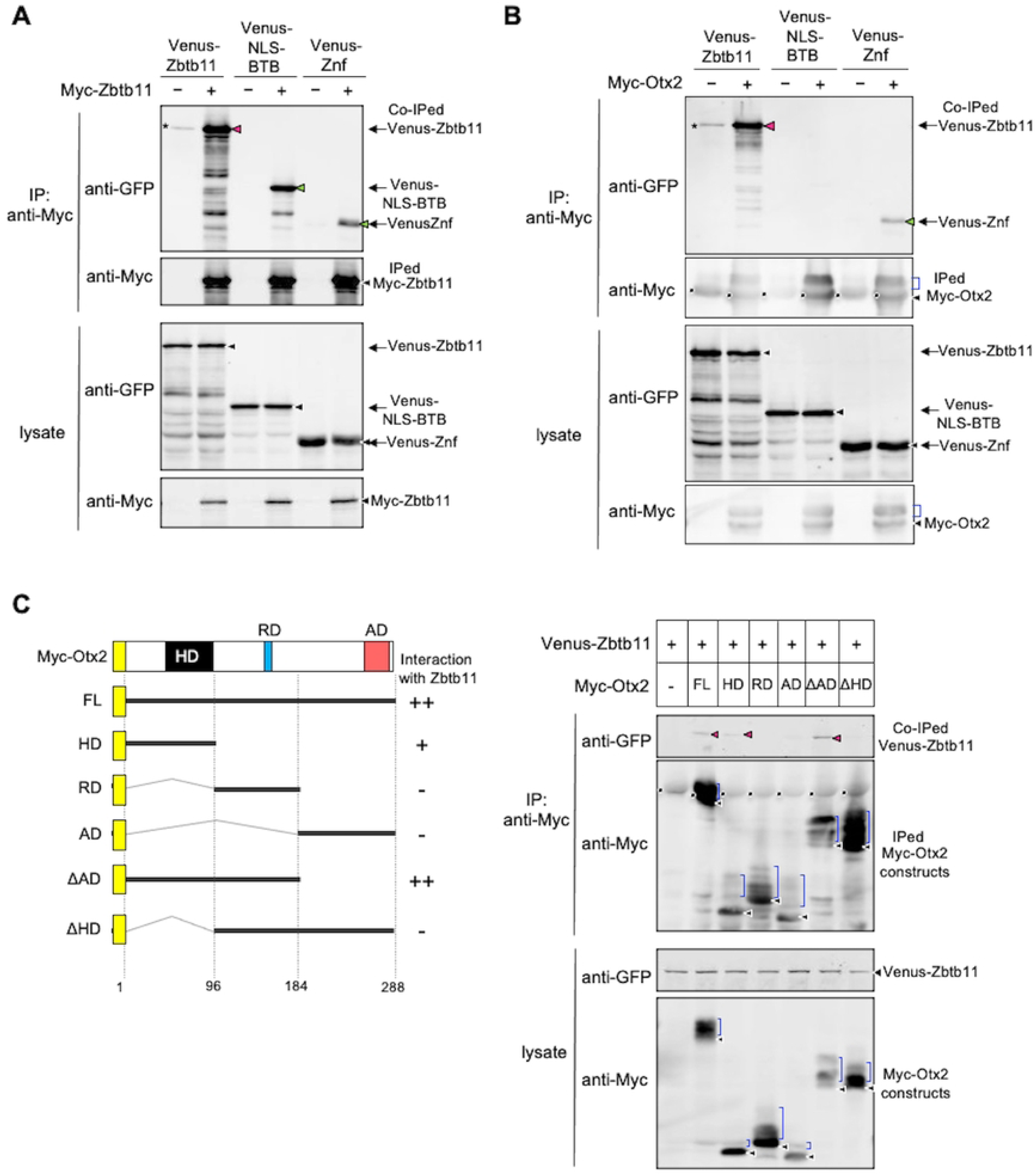
Oligomerization of Zbtb11 and the interaction between Zbtb11 and Otx2. Co-immunoprecipitation (Co-IP) assays were performed using lysates prepared at gastrula stages (stages 10.5-11), and immunoprecipitated with anti-Myc antibody. After immunoprecipitation, western blotting was performed with antibodies as indicated. The amount of protein expressed in the lysate is verified by western blotting (lysate). (A) Co-IP of Zbtb11 with its deletion constructs. mRNA for Venus-Zbtb11, Venus-NLS-BTB, or Venus-Znf was co-injected with or without mRNA for Myc-Zbtb11 into both blastomeres at the 2-cell stage. (B) Co-IP of Otx2 with the full-length or deletion constructs of Zbtb11. mRNA for Venus-Zbtb11, Venus-NLS-BTB, or Venus-Znf was co-injected with or without mRNA for Myc-Otx2. (C) The Otx2 region for interaction with Zbtb11. The left panel shows schematic structures of Myc-Otx2 and its deletion constructs. The homeodomain (HD), repression domain (RD), and activation domain (AD) are indicated. The N-terminal Myc tag is indicated as a yellow box. The full-length (FL) and deletion constructs of Otx2 are indicated by thick lines and the positions of amino acid residues are indicated. Calculated and apparent molecular masses of each construct (kDa): Myc-Otx2 FL, 43 and 52; HD, 23 and 32; RD, 21 and 34; AD, 23 and 31; ΔAD, 32 and 43; ΔHD, 32 and 41, respectively. The strength of interaction between Zbtb11 and each Otx2 construct is indicated (++, +, -) on the right side. The right panel shows the Co-IP of Zbtb11 with the full-length or deletion constructs of Otx2. mRNA for Venus-Zbtb11 was co-injected with mRNA for each Myc-Otx2 construct. Co-IPed, co-immunoprecipitated bands; IPed, immunoprecipitated bands; black arrowheads, nascent products; magenta and green arrowheads, co-immunoprecipitated Venus-Zbtb11 constructs (A-C). Asterisks indicate non-specific bands. Note that multiple bands detected below the full-length product of each Zbtb11 construct are due to partial degradation (A,B). Blue brackets, modified Myc-Otx2 constructs; black dots, IgG heavy chains from the anti-Myc antibody (B,C).

We next tested the interaction of Zbtb11 with a possible partner protein, which is expressed in the ANE and is involved in anterior patterning of the neuroectoderm. Among transcription factors expressed in the ANE, Otx2 is known to negatively regulate the expression of *gbx2* for the regionalization of the ANE [37]. Therefore, we hypothesised that Zbtb11 forms a protein complex with Otx2 for its function. To test this, we performed Co-IP assays using Venus-Zbtb11 constructs and Myc-tagged Otx2 (Myc-Otx2). The band of Venus-Zbtb11 co-immunoprecipitated with Myc-Otx2 was strongly detected (magenta arrowhead) compared to that of the negative control (asterisk) (Fig 6B), supporting their complex formation. Furthermore, deletion analysis showed that Venus-Znf but not Venus-NLS-BTB was co-immunoprecipitated with Myc-Otx2 (Fig 6B, green arrowhead), suggesting that the C-terminal region including the Znf domains of Zbtb11 is required for complex formation with Otx2. However, the co-immunoprecipitated band of Venus-Znf was weaker than that of full-length of Zbtb11, implying that the full-length is required for a full interaction with Otx2. Note that multiple bands of Myc-Otx2 indicate the presence of several phosphorylated forms that migrate more slowly than the nonphosphorylated form (Fig 6B, blue brackets and black arrowheads, respectively), as previously reported [40]. We next examined which region of Otx2 is involved in the interaction with Zbtb11 using Myc-tagged deletion constructs of Otx2 [Myc-Otx2 full-length (FL), homeodomain (HD), repression domain (RD), activation domain (AD), ΔAD, ΔHD] (Fig 6C, left). Venus-Zbtb11 co-immunoprecipitated with ΔAD as seen with FL and weakly with HD (Fig 6C, right; magenta arrowheads), but not with RD, AD, or ΔHD. This demonstrates that the homeodomain of Otx2 is required for the interaction with Zbtb11. The region containing the homeodomain and the repression domain (aa 1–184: see ΔAD) is sufficient for full interaction; i.e. RD enhances the interaction with HD (compare HD and ΔAD).

### Zbtb11 enhances repression activity of Otx2

Zbtb proteins reportedly interact with transcriptional co-repressors [11–14]. Therefore, we next tested the interaction of Zbtb11 with Tle1, a co-repressor for Otx2 [37], using Co-IP assays with Venus-Zbtb11 and HA-tagged Tle1 (HA-Tle1). As shown in Fig 7A, there was no difference in the intensity between Venus-Zbtb11 bands with or without HA-Tle1, suggesting that Zbtb11 does not interact with Tle1. Because Zbtb11 and Otx2 formed a complex (Fig 6B and 6C), we next tested whether Zbtb11 influences the interaction between Otx2 and Tle1. As shown in Fig 7B, HA-Tle1 was co-immunoprecipitated with Myc-Otx2 as previously reported [37], and the band intensity of HA-Tle1 co-immunoprecipitated with Myc-Otx2 appeared to be enhanced in the presence of Venus-Zbtb11 compared to a negative control, Venus-NLS (comparing the band in lane 2 with that in lane 5; orange arrowheads). These data suggest the possibility of tripartite interactions between Zbtb11, Otx2, and Tle1.

**Fig 7.**
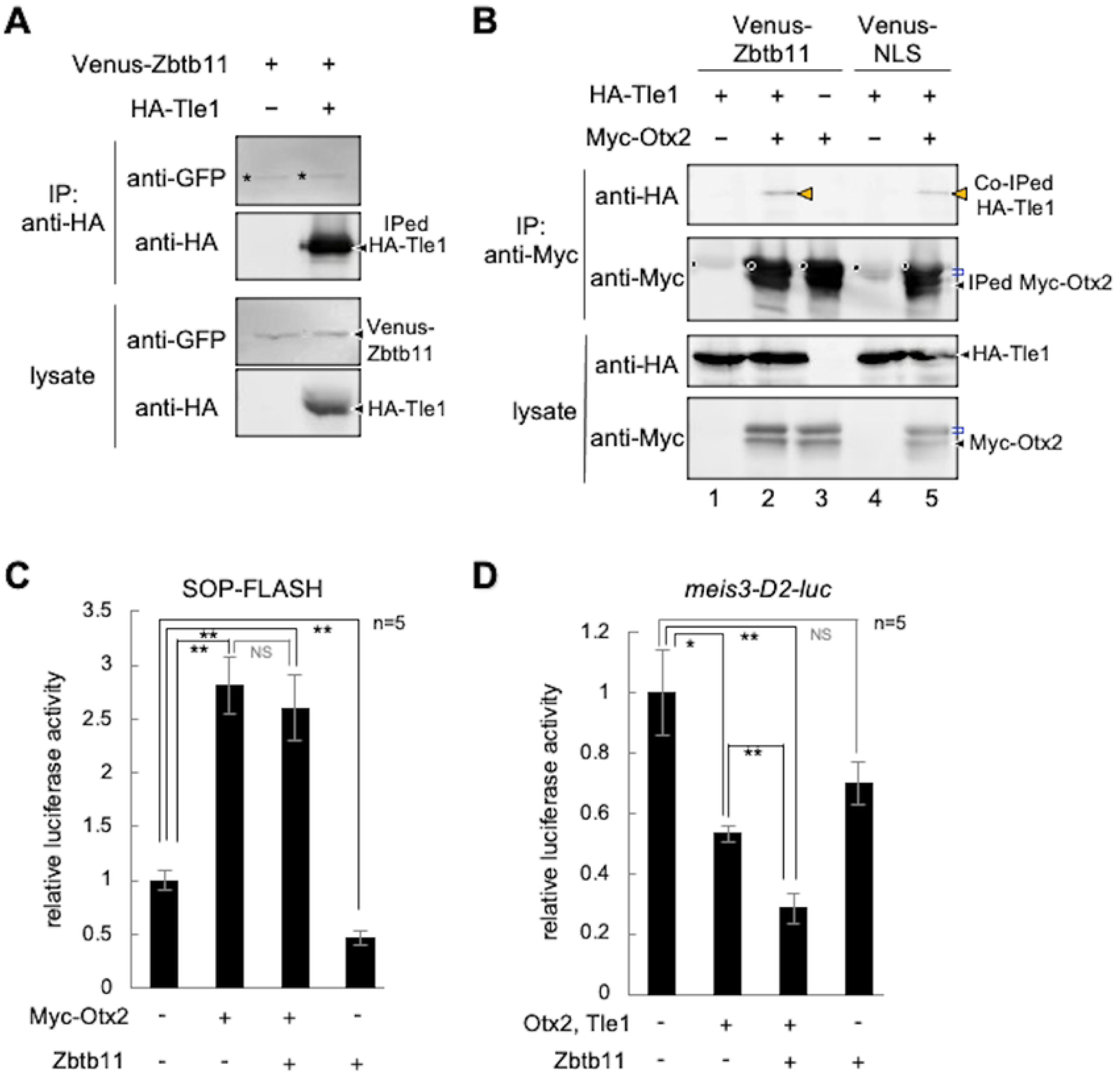
Effects of Zbtb11 on the interaction between Otx2 and Tle1. (A,B) Co-immunoprecipitation (Co-IP) assays and western blots were performed using injected embryos as described in the Fig 6 legend. (A) No specific interaction between Zbtb11 and Tle1. mRNA for Venus-Zbtb11 was co-injected with or without mRNA for HA-Tle1 as indicated. (B) Enhancement of the interaction between Otx2 and Tle1 by Zbtb11. mRNA for HA-Tle1 was co-injected with combinations of mRNA for Myc-Otx2, Venus-Zbtb11, or Venus-NLS, as indicated. The band of HA-Tle1 co-immunoprecipitated with Myc-Otx2 is increased by co-expression of Venus-Zbtb11 (lane 2), compared to that of Venus-NLS as a control (lane 5). Black arrowheads, nascent products (A,B). Asterisks indicate non-specific bands (A). Orange arrowheads, co-immunoprecipitated HA-Tle1; blue brackets, modified Myc-Otx2; black dots, IgG heavy chains from the anti-Myc antibody (B). (C) The effect of Zbtb11 on transactivation activity of Myc-Otx2 analyzed by luciferase reporter assays. mRNA for Zbtb11 with combinations of mRNA for Myc-Otx2 was co-injected with SOP-FLASH reporter DNA, as indicated. Amounts of injected mRNAs (pg/embryo): *Myc-otx2*, 100; *zbtb11*, 1500. (D) The effect of Zbtb11 on transrepression activity of Otx2 and Tle1 analyzed by luciferase reporter assays. mRNA for Zbtb11 with combinations of mRNA for Otx2 and Tle1 was co-injected with the *meis3*-D2-luc reporter DNA, as indicated. Amounts of injected mRNAs (pg/embryo): *otx2*, 20; *tle1*, 20; *zbtb11*, 1500. \**P* < 0.05, \*\**P* < 0.01 (*t*-test); error bars, standard error of the mean (s.e.m.); NS, not significant; n, the total number of samples (C,D).

We next examined whether Zbtb11 actually affects the regulation of Otx2-target genes. Because Otx2 has transactivation and transrepression activities depending on its partner proteins [39], we first tested the effect of Zbtb11 on transactivation activity of Otx2 using the luciferase reporter construct SOP-FLASH, which contains tandemly repeated *rax*-enhancer elements binding to Otx2 [38]. As shown in Fig 7C, Zbtb11 alone did not activate but rather reduced the basal level of luciferase activity, and did not significantly affect the activation of luciferase activity enhanced by Myc-Otx2. This data suggests that Zbtb11 does not affect transactivation activity of Otx2. We next tested the effect of Zbtb11 on transrepression activity of Otx2 on the posterior gene *meis3*. The *meis3-*D2-luc reporter construct, in which a silencer element of *meis3* is inserted upstream of the SV40 promoter, was repressed by Otx2 and Tle1 (Fig 7D) as reported [40]. Zbtb11 alone also showed repression activity to some extent, but co-expression of Zbtb11 with Otx2 and Tle1 further reduced the reporter activity (Fig 7D). This data suggests that Zbtb11 enhances transrepression activity of Otx2 and Tle1.

### Phosphorylation-dependent interactions between Otx2 and Zbtb11

We have previously shown that phosphorylation modifications of Otx2 confer its transrepression activity on posterior genes [40]. Therefore, the phosphorylation dependency of Otx2 was examined for physical and functional interactions with Zbtb11. Phosphorylatable serine (S) and threonine (T) residues (T115, S116, S132, and S158) of Otx2 were replaced with alanine residues for a nonphosphorylatable mutant (Otx2-4A), and with aspartate and glutamate residues for a phosphomimetic mutant (Otx2-4E) [40] (Fig 8A). Co-IP assays with HA-tagged Zbtb11 (HA-Zbtb11) and Myc-Otx2 constructs (Myc-Otx2-WT; wild type, -4A, and -4E) showed that a co-immunoprecipitated band of HA-Zbtb11 was detected with Myc-Otx2-4E and Myc-Otx2-WT (magenta arrowheads), but rarely with Myc-Otx2-4A (Fig 8B), suggesting a phosphorylation-dependent interaction between Otx2 and Zbtb11. Consistently, the reporter assay showed that repression of the *meis3*-D2-luc reporter by Otx2 WT and 4E, but not 4A was significantly enhanced by co-expression of Zbtb11 and Tle1 (Fig 8C). Likewise, repression of reporter activity by Zbtb11 and Tle1 was significantly enhanced by WT and 4E, but not 4A (Fig 8C), suggesting that cooperative repression activity of Zbtb11 and Otx2 depends on phosphorylation states of Otx2.

**Fig 8.**
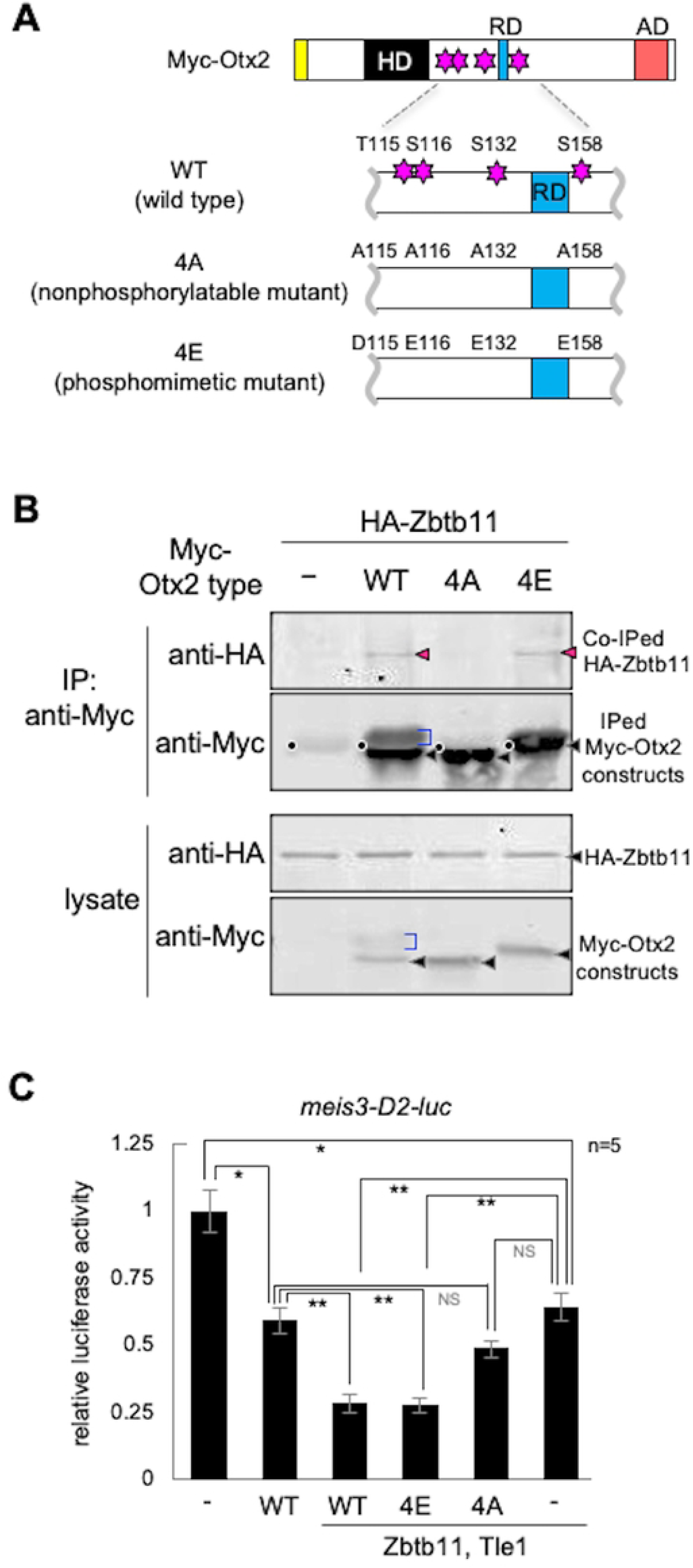
Phosphomimetic mutant of Otx2 physically and functionally interacts with Zbtb11. (A) Schematic representation of Otx2 phosphorylation sites and mutant constructs of Myc-Otx2. HD, homeodomain; RD and blue box, repression domain; AD and red box, activation domain; yellow box, Myc-tag; magenta stars, phosphorylatable serine (S) and threonine (T) residues (T115, S116, S132 and S158). Phosphorylation sites around the Otx2-RD are replaced with alanine [A] or aspartate [D] and glutamate [E] residues, as indicated. WT, wild type; 4A, nonphosphorylatable mutant; 4E, phosphomimetic mutant. (B) Co-IP of Zbtb11 and Otx2 mutants. mRNA for HA-Zbtb11 was co-injected with or without mRNA for each Myc-Otx2 construct. Black arrowheads, nascent products; magenta arrowheads, co-immunoprecipitated HA-Zbtb11; blue brackets, modified Myc-Otx2; black dots, IgG heavy chains from the anti-Myc antibody. (C) Luciferase reporter assay. mRNA for Zbtb11 with combinations of mRNA for Otx2 constructs and Tle1 was co-injected with the *meis3*-D2-luc reporter DNA, as indicated. Amounts of injected mRNAs (pg/embryo): *otx2* (*wt*, *4A*, *4E*), 20; *tle1*, 20; *zbtb11*, 1500. \**P* < 0.05, \*\**P* < 0.01 (*t*-test); error bars, standard error of the mean (s.e.m.); NS, not significant; n, the total number of samples.

## Discussion

### Repression complex of Zbtb11, Otx2, and Tle1 for anterior patterning

Based on this study and previous reports (see below), we propose a model for the role of Zbtb11 in transcriptional regulation of Otx2 in the ANE (Fig 9). This model includes the following lines of evidence: (i) Otx2 preferentially interacts with the bicoid palindromic motif, P3C (bicoid/paired-type homo- or hetero-dimer-binding motif) (TAATCNNATTA) [49,50], which is mainly required for repressive gene regulation [39,51–53]; (ii) Zbtb11 is dimerized or oligomerized through both the BTB-containing region and Znf domains (Fig 6A); (iii) Zbtb11 forms a complex with Otx2 through its Znf domains (Fig 6B); (iv) Otx2 binds to Tle1 [37], which is likely to be enhanced by Zbtb11 (Fig 7B); and (v) Zbtb11 enhances repressive activity of Otx2 for a reporter gene (Fig 7D). Thus, it is likely that dimerized or oligomerized Zbtb11 acts as a scaffold and forms a complex with two molecules of Otx2 and Tle1 for repressing posterior genes in the ANE (Fig 9, left). This model explains why loss-of-function of either Zbtb11 (this study) or Otx2 [27,54,55] causes anterior expansion of *gbx2* expression, and also why Zbtb11 gain-of-function leads to the anterior expansion of *gbx2* expression by disrupting the stoichiometric Otx2/Zbtb11 complex (Fig 9, right). Thus, the stoichiometric balance between Zbtb11 and Otx2 is required for proper repressive complex formation. However, the stoichiometry of these proteins and the actual structure of this repressive complex require further clarification.

**Fig 9.**
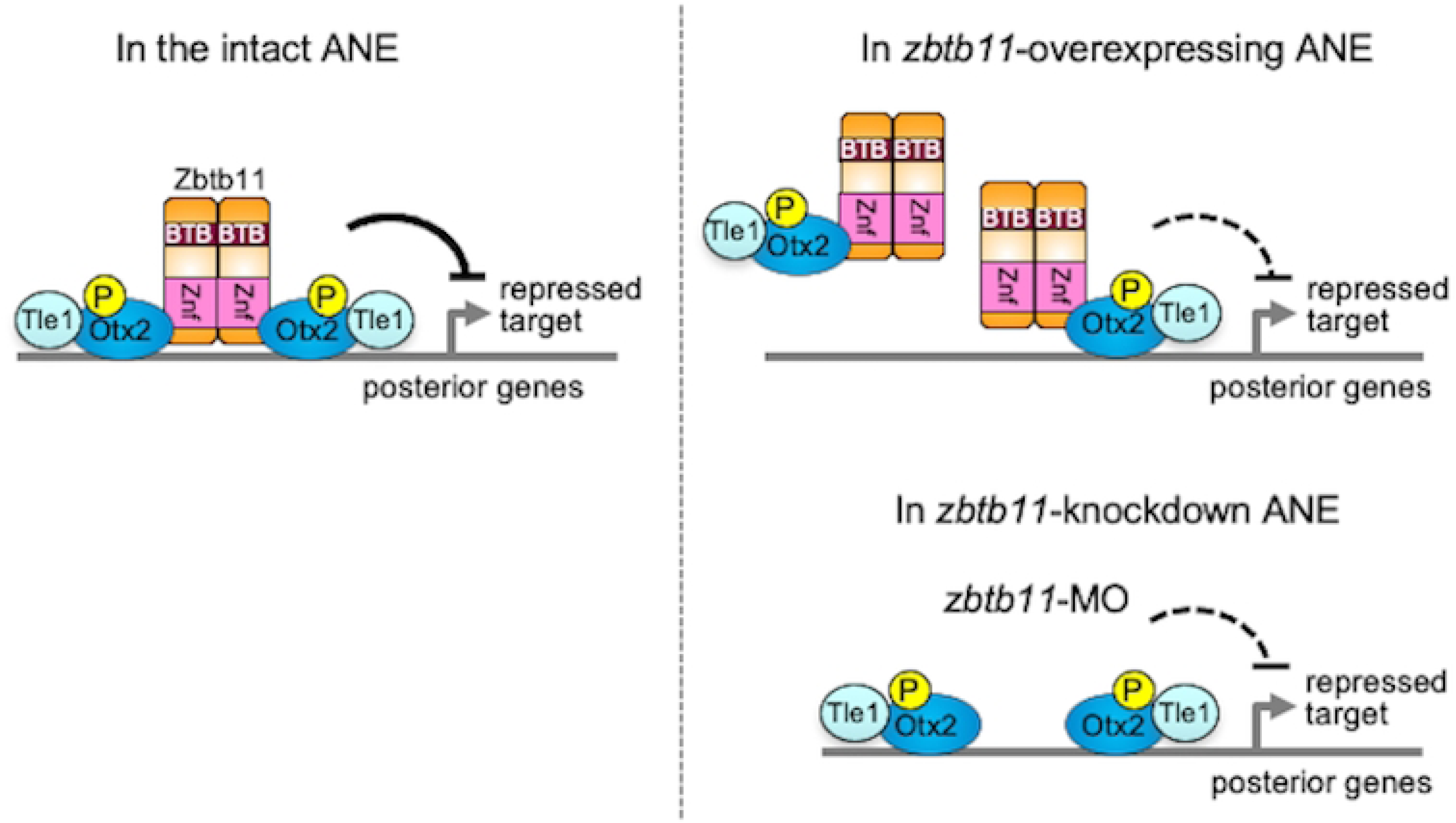
Schematic model of the complex formation between Zbtb11, Otx2 and Tle1 in the intact, *zbtb11*-overexpressing or *zbtb11*-knockdown embryo. In the intact anterior neuroectoderm (ANE), Zbtb11 forms a repressive complex with itself, phosphorylated Otx2, and Tle1 to repress posterior genes, and controls the anteroposterior patterning of the neural plate (left). However, overexpression of *zbtb11* in the ANE disrupts the stoichiometry between Zbtb11, phosphorylated Otx2, and Tle1, and prevents proper complex formation, leading to a reduction in the transrepression activity of phosphorylated Otx2 and Tle1 (right). Furthermore, knockdown of *zbtb11* by *zbtb11*-MO decreases the repression activity of phosphorylated Otx2 and Tle1, leading to derepression of posterior genes in the ANE (right).

In this study, we showed phosphorylation-dependent interactions of Otx2 with Zbtb11 in Co-IP and reporter assays (Fig 8B and 8C). Our group has previously reported that Otx2 with Gsc functions as a repressor for ventral and posterior genes in the head organizer [39]. We further reported that Otx2 requires its phosphorylation for interacting with Tle1, but not for interacting with Gsc to repress posterior genes, indicating multiple regulatory modes of Otx2-repressive complex depending on binding partners as well as cis-regulatory modules for their binding [40]. The data in this paper suggests that Zbtb11 confers the repression activity of phosphorylated Otx2 on posterior genes in the ANE (Fig 9). Thus, this study demonstrates for the first time that Zbtb11 is a repressive partner of phosphorylated Otx2.

We have shown that *zbtb11* is involved in anterior patterning of the neuroectoderm and eye formation (Fig 3 and Fig 4). In the line of the loss-of-function experiments with *zbtb11*-MO, *zbtb11* does not affect the posterior boundary of *otx2* expression even though *gbx2* expression expands anteriorly (Fig 4G). Given that *otx2*-*gbx2* mutual repression occurs in the MHB as previously reported (Fig 4H) [25–27,29], Gbx2 should repress *otx2* expression in the region where *gbx2* expands, but this is not the case. As for the regulation of *otx2* expression in the neuroectoderm, FGF8 signaling in the region posterior to the MHB is reportedly involved in the determination of the caudal limit of *otx2* expression in the midbrain [56–58]. *Fgf8* is expressed at the boundary between *otx2* and *gbx2* expression [59,60], and FGF8 signal can repress *otx2* expression and induce *gbx2* expression [56–58]. These reports imply that the posterior boundary of *otx2* is determined by *fgf8* or unknown transcription factors other than Gbx2, and *zbtb11* knockdown may not affect their expression. Regarding early eye formation, a gene cascade starting from *otx2* through *rax* to *pax6* is reported to define the eye field (Fig 4I), and this region starts to express other downstream eye-specific genes, *six3*, *lhx2*, *tll*, and *six6* [35]. Loss-of-function experiments of *zbtb11* showed that the expression of *otx2* and *rax* were not significantly affected, but that of *pax6* expanded posteriorly (Fig 4G), suggesting that *zbtb11* functions downstream of *otx2* and *rax* and upstream of *pax6* in the gene cascade of eye field specification. It also suggests that *zbtb11* delineates the posterior boundary of *pax6* expression.

### Distinct activities of the C-terminal Znf and the N-terminal BTB regions

The Znf domains of Zbtb proteins have the ability to bind to DNA in a sequence-specific manner and to mediate protein-protein interactions. For DNA binding, a recent genome-wide analysis has identified the DNA motif for Zbtb11-binding (CCGGAAG/C) in mouse embryonic stem cells and cultured cells [61]. However, there were no such Zbtb11 binding motifs in the *meis3*-D2 silencer region, implying that Zbtb11 mediates the repressor activity of Otx2 for the *meis3* gene (Fig 7D) via protein-protein interactions. An example of transcriptional regulation through the interaction with Znf domains of Zbtb proteins is PLZF (Promyelocytic leukemia zinc finger; the same as Zbtb16). PLZF interacts with a transcription factor GATA2 through its Znf region/domains and inhibits transactivation activity of GATA2 [18]. In the present study, Zbtb11 interacted with Otx2 through its Znf domains (Fig 6B) and enhanced the repressor function of Otx2 (Fig 7D), which differs from PLZF-GATA2 interactions. Notably, our data suggest that dimerization/oligomerization of Zbtb proteins require not only the N-terminal region including the BTB domain, but also the C-terminal region including Znf domains (Fig 6A). This indicates a new role for Znf domains of Zbtb proteins in self dimerization/oligomerization.

Regarding the function of the BTB domain of Zbtb proteins, the interaction with transcriptional co-repressors or histone deacetylase (HDAC) has been well documented. For example, the N-terminal BTB region of PLZF interacts with Sin3A and HDAC1 [11]; that of BCL6 (B cell lymphomas 6; the same as Zbtb27) is required for the interaction with N-CoR and SMRT [12,13]; and Kaiso (the same as Zbtb33) interacts with N-CoR through its BTB domain [14]. However, a corepressor(s) interacting with Zbtb11 is not known yet. Instead, we have shown that the Venus-NLS-BTB construct alone ectopically induced *pax6* expression (Fig 3T and 3T’). Therefore, if Zbtb11 interacts with Sin3A, N-CoR, SMRT, HDAC1, or some other repressive factors through the BTB domain, it is possible that overexpression of Zbtb11 or its N-terminal BTB region depletes those repressive factors to de-repress *pax6* expression.

### Developmental defects and disorders caused by *zbtb11* **mutations**

Several point mutants of *zbtb11* have been discovered by genetic analyses in zebrafish and humans [19,62]. The zebrafish *zbtb11* mutant *marsanne* (*mne*) has the substitution of cysteine to serine residue at position 116 (C116S) in the integrase-like HHCC motif upstream of the BTB domain (Fig 2A and S2 Fig). This mutant exhibited CNS degeneration with small eye and microcephaly-like phenotypes in addition to neutrophil deficiency [19]. In humans, two homozygous missense mutations (H729Y and H880Q in the 5th and 10th Znf motifs, respectively) are both associated with intellectual developmental disorders with microcephaly and cerebellar atrophy [62]. These mutations in ZBTB11 impair its subcellular localisation and decrease its protein stability, leading to an assumption of ZBTB11 protein dysfunction or loss of function [61,62]. Thus, the observed microcephaly in the zebrafish mutant and the genetic disorders in humans both appeared to be consistent with that in *zbtb11* morphants in *Xenopus* (Fig 4A). This implies that the zebrafish mutant *mne* exhibits loss-of-function phenotypes possibly due to its protein instability. Notably, when proteins that require proper stoichiometry for complex formation are overexpressed, they are prone to be degraded, mediated by a specific E3 ubiqutin ligase. For example, the LIM homeodomain protein Xlim1/Lhx1 forms a complex with dimerized co-factor Ldb1 in a stoichiometric manner, but Ldb1 is susceptible to the proteasomal degradation mediated by the E3 ubiqutin ligase Rnf12 when not bound to Lhx1 [63]. Furthermore, in yeast, the oligosaccharyltransferases (OST) form multiple complexes, and overexpressing OST subunits are degraded to correct the imbalanced OST subunit stoichiometry [64]. Based on this, the fact that exogenous Zbtb11 is very unstable (S3 and S4 Figs) may be explained by supposing that Zbtb11 is involved in complex formation as a scaffold in a stoichiometric manner. Importantly, it has been reported that inactive dimers or monomers of the BTB proteins, such as BCL6, ZBTB5, ZBTB18, and Kaiso, are eliminated by proteolysis via an SCF (Skp1/Cullin/F-box) E3 ligase containing FBXL17 as the F-box protein (SCF^FBXL17^) [65,66]. Crystal structural analysis revealed that wild-type or functional BTB homodimers/heterodimers bury three key degron residues for SCF ^FBXL17^, but these key degron residues of mutant or inactive BTB homodimers/heterodimers remain accessible to SCF^FBXL17^, leading to their ubiquitination and proteasomal degradation [65,66]. As Zbtb11 has two of the three conserved key degron residues, it remains to be elucidated whether stoichiometry of Zbtb11 in complex formation is regulated by a similar mechanism.

Recently, genome-wide target analysis of chromatin immunoprecipitation (ChIP) and RNA sequencing using cultured cells showed that Zbtb11 directly regulates the expression of many mitochondrial genes that are defined as intellectual disability-associated genes [61]. Furthermore, the combination of ChIP and RNA sequencing showed that Otx2 also targets genes encoding mitochondrial components, and these Otx2-bound genes are enriched in pathways, including Alzheimer’s, Huntington’s, and Parkinson’s diseases, whose common components are mitochondrial factors in the mouse developmental/juvenile cortex [67]. Our study showing the interactions between Zbtb11 and Otx2 might provide a basis for the therapeutic potential of Zbtb11 in neurodevelopmental disorders associated with mitochondrial dysfunction, as well as for elucidating the molecular mechanisms of early anterior neural development.

In summary, our study provides new insights into the molecular mechanisms of anterior patterning of the neuroectoderm by cooperation with Zbtb11 and phosphorylated form of Otx2 as well as the neurodevelopmental disorders caused by dysregulation of Zbtb11 and perhaps Otx2. It also provides a basis for post-translational regulation of Otx2 for controlling its transcriptional activities and the generalization of the molecular features of Zbtb11 to other Zbtb family proteins.

## Acknowledgements

We thank Dr. Hiroshi Mamada for cloning the 3’-portion of the *zbtb11*, and Dr. Norihiro Sudou for pCSf107_Venus_mT vector. We also thank Dr. Mariko Kondo for critical reading of the manuscript. *X. tropicalis* were obtained from the National Bioresource Project (Amphibian Research Center, Hiroshima University, Japan). The co-author of this paper, Dr. Shuji Takahashi, passed away before the submission of the final version of this manuscript. Yumeko Satou-Kobayashi accepts responsibility for the integrity and validity of the data collected and analyzed as the corresponding author.

## Author contributions

Y.S.-K. and M.T. conceived and designed the analysis. Y.S.-K. performed most experiments and analyzed the data. S.T. and Y.H. supported knockdown experiments. Y.S.-K. and M.T. wrote the manuscript. M.A. and M.T. reviewed and edited the manuscript.

## Supporting information

**S1 Fig. Developmental expression of *zbtb11* in *Xenopus laevis* and Xenopus tropicalis.**

(A) Temporal expression of *zbtb11.L* and *zbtb11.S* in *X. laevis* embryos. Expression levels (transcripts per million: TPM) are calculated from RNA-sequencing (RNA-seq) datasets of *X. laevis* developing embryos (Session et al., 2016). (B) Temporal expression of *zbtb11* in *X. tropicalis* embryos. Expression levels (transcripts ×1000) are calculated from RNA-seq datasets of *X. tropicalis* developing embryos (Owens et al., 2016). Images are generated using Xenbase (http://www.xenbase.org/) and developmental stages (oocyte and Nieuwkoop-Faber [NF] stages) are as indicated (A,B).

**S2 Fig. Alignment of Zbtb11 amino acid sequences between different species.**

Alignment of Zbtb11 amino acid sequences between *Xenopus laevis* (Xl), *Xenopus tropicalis* (Xt), *Homo sapiens* (Hs), *Mus musculus* (Mm), and *Danio rerio* (Dr). *Xenopus laevis* has two homeologs: L and S genes. Light blue boxes, the conserved regions CR1, CR2 and CR3; purple box, the integrase-like histidine-histidine-cysteine-cysteine (HHCC) motif; brown box, the BTB domain; magenta boxes, C2H2 type zinc fingers. The blue-coloured cysteine residue (C116) indicates a mutation site in the neutrophil-deficient zebrafish mutant (marsanne, mne). Red histidine residues (H729 and H880) indicate ZBTB11 mutation sites that are associated with intellectual disability. Arrows indicate the end of the Venus-BTB construct, and the beginning of the Venus-Znf construct.

**S3 Fig. Stability of Myc-Zbtb11, Venus-Zbtb11, and its deletion constructs in the *Xenopus laevis* embryo.**

mRNA for Myc-Zbtb11, Venus-Zbtb11, or its deletion constructs was injected into two blastomeres at the 2-cell stage. Lysates were prepared at stages 10–10.5, and western blotting was performed with antibodies as indicated. (A) Western blotting of Myc-Zbtb11. SDS-polyacrylamide gel electrophoresis (PAGE) was carried out with a 7.5% polyacrylamide gel. (B) Western blotting of Venus, Venus-BTB, Venus-Znf, and Venus-Zbtb11. SDS-PAGE was carried out with a 10% polyacrylamide gel. Black arrowheads, nasent proteins (undegraded products). Calculated molecular weights (MW) of Zbtb11 constructs are as indicated besides dotted lines: Myc-Zbtb11, 140; Venus-Zbtb11, 160; Venus-BTB, 91; Venus-Znf, 87; Venus, 30. Brackets indicate degradation products, and the corresponding degradation products in the different constructs are indicated by the same colour (orange, magenta, or blue) and number of dots. White dots indicate non-specific bands. Molecular masses of protein size marker (kDa) are indicated on the right side. Uninj., uninjected sample.

**S4 Fig. Protein levels of Venus-Zbtb11 expressed in embryos.**

(A) Fluorescence observation of Venus-NLS and Venus-Zbtb11. mRNA for Venus-NLS or Venus-Zbtb11 was injected into any one of the blastomeres at the 4-cell stage and fluorescence was observed at stages 14, 19–20, and 28–30. There are no differences in the fluorescence intensity between injected regions, indicating that the protein stability of Venus-Zbtb11 does not depend on tissue type. Merged, fluorescence images merged with bright-field images; Venus, fluorescence images. (B,C) Comparison of protein expression levels between Venus-NLS and Venus-Zbtb11. mRNA for Venus-NLS or Venus-Zbtb11 was injected into both blastomeres at the 2-cell stage. Lysates were prepared at stages 10, 16, and 19 and subjected to western blotting using anti-GFP and anti-*β*-tubulin antibodies. (B) Western blotting of Venus-NLS and Venus-Zbtb11. Upper panels, bands of Venus-NLS and Venus-Zbtb11; lower panels, bands of *β*-tubulin as a loading control. (C) Expression levels of Venus-NLS and Venus-Zbtb11. The relative expression level was obtained by dividing the band intensity of Venus-NLS or Venus-Zbtb11 by that of *β*-tubulin. Stages (st.) are indicated, respectively.

**S5 Fig. Specificity of *zbtb11*-morpholino oligo (MO).**

mRNA for *zbtb11*-MO target sequences fused with eGFP (Zbtb11-ATG-eGFP construct) was injected into two blastomeres at the 2-cell stage and then *zbtb11*- MO or control MO was co-injected into all blastomeres at the 4-cell stage. Lysates were prepared from gastrula embryos and subjected to western blotting with anti-GFP and anti-*β*-tubulin antibodies. Translation of *zbtb11*-eGFP mRNA was blocked by *zbtb11*-MO injection compared to control MO-injected or uninjected samples. Amounts of injected mRNA for Zbtb11-ATG-eGFP (pg/embryo), 500; injected MOs (pmol/embryo), 1.

**S6 Fig. Original, uncropped and minimally adjusted images of western blots.**

Original images for (A) Fig 6A-C, (B) Fig 7A and 7B, (C) Fig 8B, (D) S3 Fig, (E) S4B Fig, and (F) S5 Fig are shown. See the corresponding figure legends for details. Boxes indicate the cropped images presented in the figures. M, protein size markers. Molecular masses (kDa) of the protein size markers are as indicated.

**S1 Table. The list of plasmid constructs.**

**S2 Table. The list of cutting sites and RNA polymerases for the in vitro transcription of anti-sense RNA probe.**

